# Exploring memory-related network via dorsal hippocampus suppression

**DOI:** 10.1101/2024.06.03.597201

**Authors:** Xu Han, Samuel R. Cramer, Dennis C.Y. Chan, Nanyin Zhang

## Abstract

Memory is a complex brain process that requires coordinated activities in a large-scale brain network. However, the relationship between coordinated brain network activities and memory-related behavior is not well understood. In this study, we investigated this issue by suppressing the activity in the dorsal hippocampus (dHP) using chemogenetics and measuring the corresponding changes in brain-wide resting-state functional connectivity (RSFC) and memory behavior in awake rats. We identified an extended brain network contributing to the performance in a spatial-memory related task. Our results were cross-validated using two different chemogenetic actuators, clozapine (CLZ) and clozapine-N-oxide (CNO). This study provides a brain network interpretation of memory performance, indicating that memory is associated with coordinated brain-wide neural activities.

**Significance Statement:** Successful memory processes require coordinated activity in a large-scale brain network, extending beyond a few key, well-known brain regions like the hippocampus. However, the specific brain regions involved and how they orchestrate their activity that is pertinent to memory processing remain unclear. Our study, using a chemogenetics-rsfMRI- behavior approach in awake rats, elucidates a comprehensive framework of the extended memory-associated network. This knowledge offers a broader interpretation of memory processes, enhancing our understanding of the neural mechanisms behind memory function, particularly from a network perspective.

## Introduction

Memory is a complex mental process in the mammalian brain. Extensive research has been carried out to identify individual brain regions involved in memory, such as the hippocampus, thalamus and prefrontal cortex (Murray et al., 2007; Euston et al., 2012; Danet et al., 2015; Hainmueller & Bartos, 2020). In addition, successful memory processes require coordinated large-scale communication between these key brain regions. (Aggleton et al., 2010; Rajasethupathy et al., 2015; Opalka & Wang, 2020). However, how brain regions in the memory-related network orchestrate their activity for memory process remains elusive.

To begin unraveling the relationship between brain network activity and memory performance, one research strategy involves the intentional perturbation of a brain node in the memory network, along with subsequent measurement of the corresponding changes in network activity/connectivity that covary with perturbation-induced memory changes at the behavioral level. For this purpose, the dorsal hippocampus (dHP) is an ideal candidate region because of its pivotal role in multiple memory processes: encoding, storage, and retrieval (Restivo et al., 2015; Park et al., 2016; Hainmueller & Bartos, 2020; Liu et al., 2012). Additionally, spatial and object information are encoded in the dHP as neural ensembles (Hainmueller & Bartos, 2018; Danielson et al., 2016). Anatomically, dHP is composed of dorsal regions of the dentate gyrus, CA3, and CA1. It receives input from the entorhinal cortex (EC) via the perforant pathway, which is in turn connected with the rest of the cortex (Sewards & Sewards, 2003; Witter et al., 2017). The dHP, through complex neural pathways mediated by the fornix, communicates with various brain structures such as the thalamus, mammillary complex as well as cortex (Aggleton et al., 2022). These interactions facilitate the hippocampus’ role in memory and spatial navigation. Given this extensive connectivity structure that might be implicated in memory process, a perturbation to the dHP could impart a widespread effect to other regions in the memory-related network.

In order to provide a targeted, local perturbation of the dHP, here we employed a chemogenetic approach in rats. In combination with neuroimaging techniques and behavioral assessment, we aimed to establish a relationship between memory network dynamics and memory performance. More specifically, neural activity of the dHP was suppressed using the <u>d</u>esigner receptors <u>e</u>xclusively <u>a</u>ctivated by <u>d</u>esigner <u>d</u>rugs (DREADD) technique (Armbruster et al., 2007). Resting state functional magnetic resonance imaging (rsfMRI) experiments were conducted to measure brain-wide changes in the spontaneous blood oxygen level dependent (BOLD) signal and resting state functional connectivity (RSFC) following acute DREADD-induced suppression. On separate days from imaging, spatial memory behavioral testing (Y maze) (Conrad et al., 1996; Jung et al., 2008) also took place after DREADD-induced suppression to investigate relationships between neural network changes and the corresponding changes in memory performance. rsfMRI data was acquired without the use of anesthesia, permitting a linkage between the imaging and behavioral data (Dopfel & Zhang, 2018; Tu et al., 2021; Beloate & Zhang, 2022). Moreover, as DREADD actuators can interact with multiple and diverse endogenous receptors (Martinez et al., 2019; Jendryka et al., 2019; Nagai et al., 2020), we separately utilized two DREADD actuators (CLZ and CNO) across all experiments to control for possible off-target effects. Our results show that dHP suppression causes brain-wide alterations of RSFC that are correlated with spatial memory performance. This data offers new insights into structures of the memory network that may be necessary for proper memory function.

## Methods and Materials

### Animals

Data were collected from 36 male Long-Evans rats weighing between 300-400 g. Rats were singly housed post-surgeries with food and water provided *ad libitum*. The housing room was maintained at constant room temperature and adhered to a 12-hour light and 12-hour dark cycle. All experiments in the current study were approved by the Institutional Animal Care and Use Committee (IACUC) at the Pennsylvania State University.

### Surgery

The rat brain was injected with adeno-associated viral vectors (AAVs) by way of aseptic stereotaxic surgery. Following a brief induction of isoflurane anesthesia, an intramuscular injection of ketamine (40 mg/kg) and xylazine (12 mg/kg) was administered to maintain the anesthetic state during surgery. Buprenorphine ER-LAB (1 mg/kg) was subcutaneously administered to provide post-surgery analgesia. Dexamethasone (0.5 mg/kg) and Enrofloxacin (2.5 mg/kg) were subcutaneously administered as an anti-inflammatory drug and antibiotic, respectively. Bupivacaine (4 mg/kg) was injected under the scalp for local analgesia. Heart rate and SpO2 were continuously monitored using a pulse oximeter (MouseSTAT Jr., Kent Scientific Corporation) during surgery while a heating pad (PhysioSuite, Kent Scientific Corporation) placed underneath the rat maintained the body temperature at ∼37 °C. Craniotomies were made on the skull for viral injection. For animals in the DREADD groups, AAV8-hSyn-hM4Di-mCherry (1μL per site, titer ≥ 3 ×10^12^ vg/mL, Addgene) was stereotactically injected into the dHP (coordinates: AP −3.5 mm, ML ± 2 mm, DV − 2.5 to − 2.9 mm). 13 rats received unilateral injection in the right dHP (termed unilateral DREADD group hereafter), and 15 rats received bilateral injection in the dHP (termed bilateral DREADD group hereafter). For the sham group (n = 8), AAV8-hSyn-EGFP (1μL per site, titer ≥ 3 ×10^12^ vg/mL, Addgene) was injected bilaterally in the dHP. Bone wax (Ethicon Inc.) was used to prevent bleeding from the burr holes. Animals were given at least four weeks for full recovery and sufficient DREADD protein expression (Tu et al., 2021).

### Actuator injections

Two DREADD actuators were separately used in this study: clozapine (CLZ, Sigma-Aldrich) and Clozapine N-Oxide (CNO, Sigma-Aldrich). CLZ was dissolved in DMSO and 0.1 N HCl and then diluted with saline to the concentration of 0.1 mg/ml. CNO was dissolved in DMSO and then diluted with saline to the concentration of 1 mg/ml. Control vehicles were saline solution with 0.5% DMSO. Across all experiments, the CLZ (0.1 mg/kg), CNO (1 mg/kg), and saline injections were administered intraperitoneally (IP) while animals were anesthetized with isoflurane vapor. For all behavior tests, isoflurane exposure was brief and immediately discontinued following IP injection. For saline and CLZ, animals were given at least 10 minutes to wake up before behavior tests commenced. In accordance with the literature, at least 30 minutes were given for behavior tests involving CNO injections (Jendryka et al., 2019). For imaging experiments, due to subject set up and prerequisite imaging sequences, at least 30 minutes elapsed between IP injection and rsfMRI data acquisition.

### fMRI experiment

To minimize animals’ stress and motion during awake fMRI imaging, rats were habituated to the restraining and fMRI scanning environment for 7 days (15, 30, 45, 60, 60, 60 and 60 min/day, respectively). Detailed acclimation procedures can be found in our previous publications (Tu et al., 2021; Han et al., 2022). All fMRI data were acquired on a 7T Bruker BioSpec 70/30 scanner interfaced with ParaVision 6.0.1 (Bruker, Billerica, MA). Prior to imaging, rats received IP injections with either saline, CLZ or CNO. T2*-weighted gradient-echo images covering the whole brain were acquired with an echo planner imaging (EPI) sequence using the following parameters: flip angle = 60°, repetition time (TR) = 1000 ms, echo time (TE) = 15 ms, slice thickness = 1 mm, slice number = 20, field of view (FOV) = 3.2 x 3.2 cm^2^, in-plane matrix size = 64 x 64. 600 volumes were collected for each fMRI scan, and three scans were collected for each fMRI session. A given animal was subjected to at least 3 fMRI sessions, allowing for each rat to have at least one session with each of the 3 types of injection solutions (saline, CLZ, or CNO), which were randomly ordered.

### fMRI data preprocessing

The fMRI signal preprocessing used an in-lab developed pipeline detailed in (Ma & Zhang, 2018; Liu et al., 2020). The first 10 volumes were removed to ensure a steady state of MR signal. Framewise displacement (FD) was calculated to estimate the motion level of each fMRI volume. Volumes with FD > 0.5 mm were discarded together with their 2 previous and 5 following volumes. Volumes with FD in the range of 0.3-0.5 mm were discarded with 1 previous and 3 following volumes. Volumes with FD in the range of 0.2-0.3 mm were discarded with their immediate neighboring volumes. Scans that had over 20% of volumes removed were excluded from further analysis. Motion-scrubbed fMRI images were registered to a predefined T2-weighted image template through rigid body transformation. Registered images were motion corrected using SPM12 package, and distortions of the images were corrected using the Advanced Normalization Tools (ANTs, Avants et al., 2008). Independent component analysis (ICA) (Calhoun et al., 2001) was applied to identify non-neural artifacts. Time courses of noise IC components, motion parameters, and signals from the white matter (WM) and cerebrospinal fluid (CSF) were voxel-wise regressed out from the fMRI signal. fMRI data were then spatially smoothed (FWHM = 1 mm). Finally, for RSFC analysis, fMRI data were temporally bandpass (0.01- 0.1 Hz) filtered and truncated to the first 480 volumes.

### fMRI data postprocessing

Fractional amplitude of low-frequency fluctuations (fALFF) was voxel-wise calculated by normalizing the signal amplitude in the low-frequency range (0.01-0.08 Hz) to the signal amplitude in the full frequency range (Zou et al., 2008). This quantity was used to estimate the BOLD amplitude at the resting state. For RSFC analysis, the whole brain was parcellated into 65 bilateral ROIs using the Swanson atlas (Swanson, 2004). RSFC between pairwise ROIs was calculated using the Pearson’s correlation of their BOLD time series. To generate group level results for z-transformed RSFC, we employed multiple linear mixed effects (LME) models. Firstly, to control for off-target effects of CNO and CLZ in RSFC analysis, sham control data were analyzed using the following LME model:

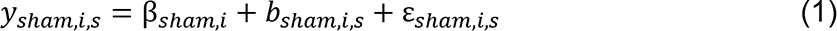

Here, *y*_*sham,i,s*_ represents observation *i* of sham subject *s*, where each observation represents a scan-derived parameter (i.e. RSFC value for a given pairwise ROI) from individual saline, CNO, or CLZ scans in sham rats. β_*sham,i*_ represents the grand mean of the whole-model average, *b*_*sham,i,s*_ is the subject-based mean difference with the grand mean for sham subject s, and ε_*sham,i,s*_ is the observation error for each scan of the subject. After model approximation, the estimated β_*sham,i,saline*_ was subtracted from both the β_*sham,i,CNO*_ and β_*sham,i,CLZ*_ estimates. The resulting ‘sham difference in RSFC’ was then subtracted from the corresponding DREADD data sets to account for any suprathreshold or subthreshold effects of each actuator. Once this correction was applied (measured as β_0_,*corrected,i* in DREADD datasets), each group (sham/unilateral/bilateral) was then modeled separately. For each DREADD group, the following model was used:

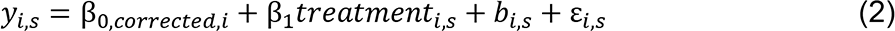

In the sham group, the following model was used:

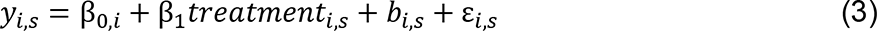

where *treatment*_*i,s*_ was modeled separately and categorically represented as 0 for saline condition and 1 in CNO/CLZ conditions. After algorithmic estimation of all coefficients, the significance of β_1_was calculated against the null hypothesis of β_1_ = 0 . The model compared saline scans to CNO and CLZ scans, respectively, to assess whether the given actuator produced a statistically significant effect on z-transformed RSFC values for a particular group. Matlab function ‘fitlme’ was used for the analysis.

### Locomotion test

After all imaging experiments, each rat was tested in three locomotion-assessment sessions with either saline, CLZ (0.1 mg/kg) or CNO (1 mg/kg) injection; all rats received all injection types in a randomized fashion. A given rat’s sessions were separated by at least 7 days. Prior to each test, rats were habituated to the testing room for 15 minutes in their home cages. In each session, following the injection of saline/CLZ/CNO, home cage locomotion was recorded by an infrared camera for 30 minutes. Locomotor parameters including total head distance travelled and total body distance travelled were analyzed by ANY-maze (Stoelting. Co).

### Y maze

After all locomotion experiments, spatial memory was examined using two-trial Y-maze tests (Conrad et al., 1996). Like locomotion tests, each rat was tested in all three different treatment conditions (saline/CLZ/CNO), which were randomly ordered and separated by at least 7 days.

The Y-maze was composed of three identical arms (50×16×32 cm^3^) made of opaque black acrylic plastic plates with a 120° angle separating neighboring arms (Conrad et al., 1996). The three arms are labeled with distinct names: Start Arm, Novel Arm, and Other Arm. There were no visual cues inside the arms. However, outside the boundaries of the Y-maze, colored paper and lab equipment with different shapes could be used as visual cues, which were rearranged across different treatment conditions (saline/CLZ/CNO) for each rat. For each subject, the order of visual cue presentation was randomized across treatment condition.

Each Y-maze test was composed of two trials. In Trial 1, an opaque black plastic plate was used to block the Novel Arm. After the injection of saline/CLZ/CNO, the rat was placed into the Start Arm and allowed to explore the Y-maze for 15 minutes. The Start Arm, Novel Arm, and Other Arm were randomly assigned for each test but remained fixed between the two trials. At the end of Trial 1, the rat was returned to its homecage outside of the testing room for a fixed inter-trial interval (ITI = 2 hours). After the ITI elapsed, Trial 2 then began by placing the rat in the Start Arm; this time, though, all arms were available for exploration. The rat was allowed to explore all 3 arms for 5 minutes. The movement of the rat was recorded by an infrared camera fixed directly above the Y-maze. The *first* new arm entered (the Novel arm or the Other arm) and total time spent in each arm were analyzed by ANY-maze (Stoelting. Co). 2×2 Chi-square tests were applied to compare the number of first entries to the Novel arm between saline- and actuator-treated conditions. Percentage of total time spent in each arm was calculated for each rat and compared between saline and each actuator-treated condition using paired t-tests.

### Correlating Y-maze Data to RSFC Data

In order to establish a relationship between Y-maze outcomes and RSFC data, a correlative analysis was carried out for each group (sham/unilateral/bilateral). Within each group, for a given injection (saline/CNO/CLZ), the RSFC between a given ROI pair A, B, measured by the z-transformed Pearson’s correlation (*r*) of their rsfMRI time series was averaged across an individual subject’s scans. The average correlation coefficient (*r̄*) produced from the saline scans was then subtracted from that of both CNO and CLZ to produce Δ*r̄*. For the same subject, the Y-maze novel arm time spent in the saline condition (*t*) was subtracted from the novel arm time spent in the corresponding actuator condition (CNO/CLZ) to produce Δ*t*. This process was repeated across all *n* subjects in the group generating a collection of values of the form: (Δ*t*_1_, Δ*t*_2_, ⋯, Δ*t*_n_) and (Δ*t*_1_, Δ*t*_2_, ⋯, Δ*t*_n_). Subsequently, Pearson’s correlation was calculated between the two lists to generate a correlation coefficient *P*_*A,B*_. This process was completed across all ROI pairs, resulting in a group-specific, actuator-specific matrix of correlation coefficients *P*.

### Electrophysiology

At the conclusion of all imaging and behavior experiments, a subset of animals was subjected to non-survival electrophysiology to validate viral expression functionality. Electrophysiology was recorded in 4 sham rats and 4 bilateral DREADD rats in the anesthetized condition. Rats were initially anesthetized using 3-4% isoflurane and xylazine (12 mg/kg). The isoflurane concentration was then lowered to 0.5% and maintained at this level. In this condition, a single-channel electrode (NeuroNexus) was gradually inserted into the dHP (AP −3.5 mm, ML 2 mm, DV − 2.7 mm). A reference wire was attached to the skull tissue, and a grounding wire was connected to the stereotaxic frame. Following the initiation of isoflurane administration, at least 40 minutes elapsed before the experiment proceeded; heart rate and electrophysiological signal were monitored to ensure that the animal reached a stable, lightly anesthetized state. Subsequently, the animal received an injection of saline, and the electrophysiological signal was recorded for 20 minutes as a baseline measurement. After 20 minutes of baseline recording, 0.1 mg/kg CLZ was injected intraperitoneally; electrophysiology recordings continued for another 40 minutes. The total length of recording (NeuroNexus, sampling rate = 20 kHz) was 60 min. CNO was not evaluated because appropriate responses to CLZ would necessarily imply appropriate responses to CNO (hM4Di is responsive to both actuators).

Raw data were notch filtered at 60 Hz to filter out power line noise. Multi-unit activity (MUA) signal was obtained by high-pass filtering the data at 1000 Hz. Individual points of the recorded signals were classified as spikes if signal amplitude exceeded 4 times the standard deviation of the entire high-passed filtered signal, with a minimal window of 10 milliseconds following the previous spike-classified point (Tu et al., 2021).

### Histology

Histology was carried out following the completion of all experiments in all animals to confirm the location of DREADD protein expression (Fig. S1). Animals were first anesthetized and then perfused with saline and 4% paraformaldehyde (PFA) solution. The brains were stored in 4% PFA and 30% sucrose solution for a week and cut into 60 μm thick slices. Fluorescence signal was imaged using a Leica SP8 DIVE multiphoton microscope.

## Results

AAVs expressing inhibitory G-protein coupled hM4Di receptors with a pan-neuronal synapsin promoter (AAV8-hSyn-hM4Di-mCherry) were stereotactically injected into the bilateral (bilateral DREADD group, n = 15) or unilateral (right hemisphere, unilateral DREADD group, n = 13) dHP. Fig. 1 shows the DREADD protein expressed in regions that coincide with dorsal DG and CA3, prominent subregions of the dHP. Control null virus vectors (AAV8-hSyn-EGFP) were injected into the bilateral dHP in sham animals (n = 8). Animals were given at least 4 weeks for recovery and protein expression. Two DREADD actuators, CLZ (0.1 mg/kg) and CNO (1 mg/kg), were used in different sessions for fMRI and behavior experiments to account for potential off-target effects, as suggested by a previous study (Jendryka et al., 2019).

**Figure 1.**
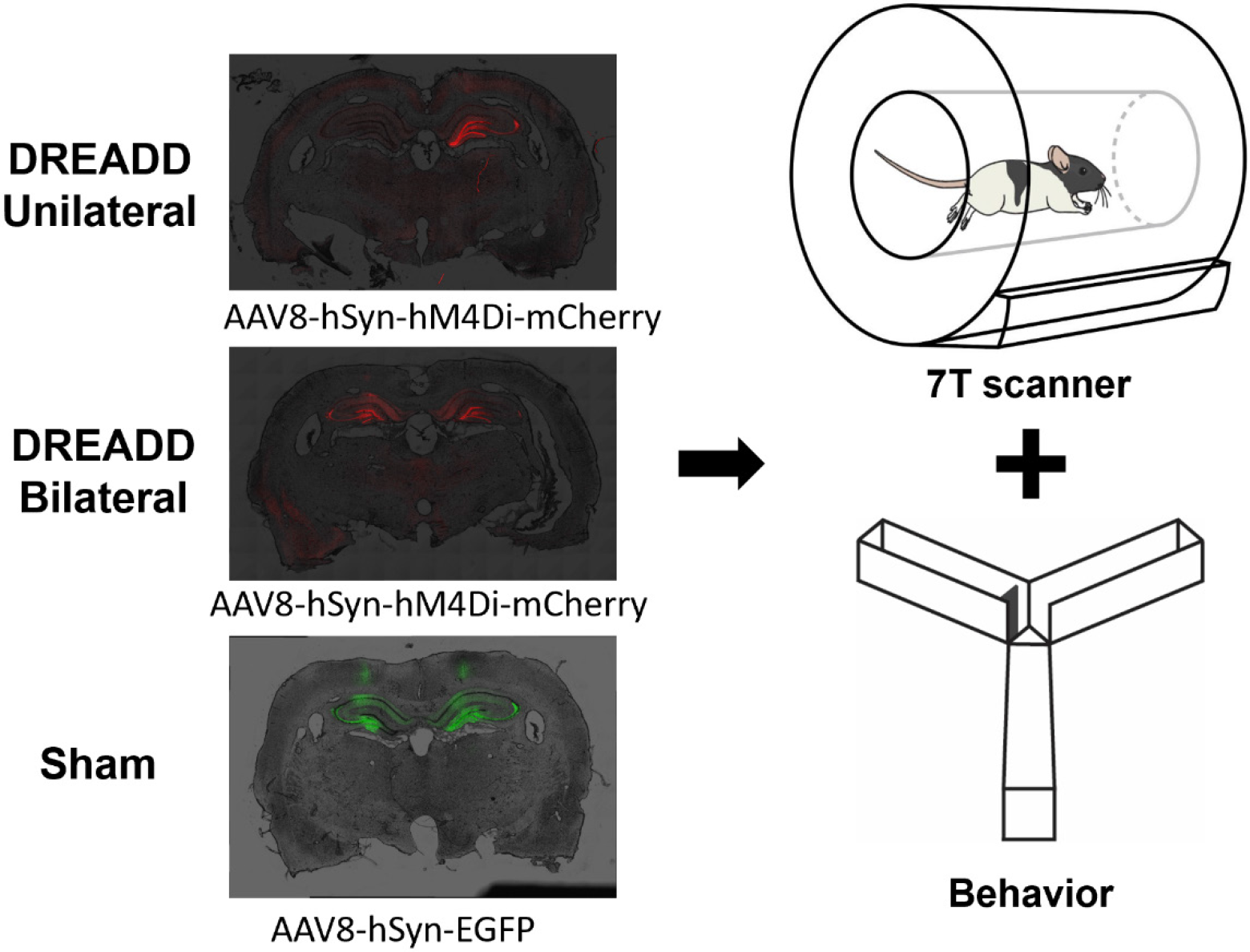
Schematic diagram of the experimental setup and paradigm. hM4Di- expressing AAVs were injected in either the unilateral or bilateral dorsal hippocampus (dHP) of the rat brain. The sham group received bilateral-dHP injection of EGFP-expressing AAVs. Rats were subjected to rsfMRI imaging and Y-maze behavioral test.

### hM4Di DREADDs suppress dHP neural activity

To validate the neuronal suppression effect of hM4Di-DREADDs, we recorded the electrophysiological signal in the dHP before and after actuator administration in a lightly anesthetized state (isoflurane 0.5%, sham rats, n = 4; bilateral DREADD rats, n = 4). After the animal reached a stable anesthetized state, the dHP electrophysiological signal was first recorded for 20 minutes as a baseline measurement. Subsequently, 0.1 mg/kg CLZ was injected intraperitoneally as the electrophysiology recording continued for another 40 minutes. Raw data of neural recording show that the firing rate was markedly reduced after CLZ administration in DREADD rats (Fig. 2a). Two-way mixed-effect ANOVAs with rat group (DREADD vs sham) and time (baseline vs post-CLZ window) as between and within factors, respectively, revealed that the firing rate was significantly reduced from 10 minutes after CLZ injection in DREADD animals (10-20 min window, p_interaction_ = 0.015) and the suppression effect persisted throughout the recording session (20-30 min, p_interaction_ = 0.0055; 30-40 min, p_interaction_ = 0.0042) (Fig. 2b). In contrast, sham rats did not show significant changes in the firing rate (paired t-tests: baseline vs 0-10 min, p = 0.70; baseline vs 10-20 min, 0.68; baseline vs 20-30 min, p = 0.40; baseline vs 30-40 min, p = 0.28, Fig. 2b). These data confirmed the neuronal suppression effect of hM4Di-DREADDs in the dHP and null effect of CLZ alone in sham rats.

**Figure 2.**
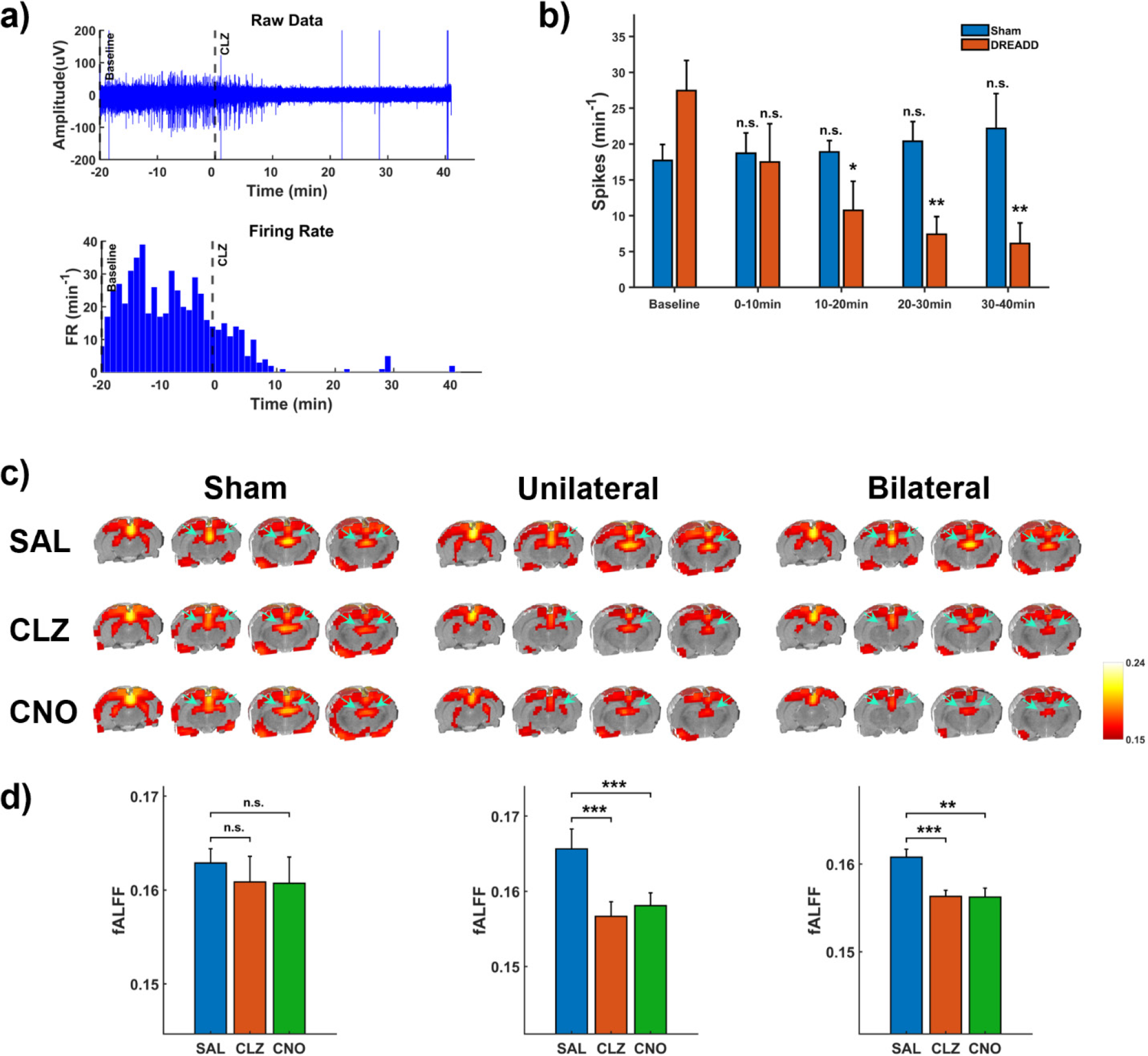
Validation of neuronal suppressing effect of hM4Di-DREADDs. a) Representative electrophysiological signal in a bilaterally hM4Di-DREADD expressed rat. Upper: Raw electrophysiological data. lower: Firing rate per minute. **b)** Statistical analysis of the actuator effect on the spiking rate in sham (n = 4) and DREADD (n = 4) rats. The effect of actuator in sham rats was evaluated by multiple paired t-tests (baseline vs 0-10 min, p = 0.70; baseline vs 10-20 min, p = 0.68; baseline vs 20-30 min, p = 0.40; baseline vs 30-40 min, p = 0.28). Actuator effect in DREADD rats was analyzed using multiple two-way mixed-effect ANOVAs with the rat group (DREADD vs Sham) and time (baseline vs post-actuator window) as between and within factors, respectively. The spiking rate was averaged over the 20 minutes of baseline and averaged over each 10-min window following CLZ administration (0 - 10 min, pinteraction = 0.077;10 - 20 min, pinteraction = 0.015; 20 - 30 min, pinteraction = 0.0055; 30 - 40min, pinteraction = 0.0042). **c)** Group-level raw fALFF maps across different injection conditions. Arrows indicate the location of the dHP. **d)**

The suppression of dHP activity by hM4Di-DREADDs was also confirmed based on the rsfMRI data. We first confirmed the actuators do not have biased influence on motion or respiration rates, as these factors could alter the rsfMRI signal (Fig. S2) (Power et al., 2014; Teichert et al., 2010). Subsequently, the activity of the dHP was quantified by fALFF, which measures the amplitude of spontaneous BOLD activity by calculating the ratio of signal amplitude in the low-frequency (0.01-0.08 Hz) range to the total signal amplitude in the full spectrum (Zou et al., 2008). Fig. 2c shows the raw voxel-wise fALFF values in sham, unilateral, and bilateral DREADD rats after the administration of saline, CLZ or CNO. In Fig. 2d, voxel-wise fALFF values were averaged throughout the dHP, and the effect of the actuators was assessed by LME models. In sham rats, no statistically significant difference of fALFF in the dHP was observed between saline and either actuator-treated condition (t-tests: SAL vs CLZ, p = 0.62; SAL vs CNO, p = 0.60; Fig. 2d). Both CLZ and CNO treatments significantly decreased the fALFF in the dHP of unilateral and bilateral DREADD animals, as confirmed by two-way ANOVAs with virus type and treatment as main factors (unilateral DREADD rats: CLZ treatment, p_interaction_ = 4.09×10^-4^; CNO treatment, p_interaction_ = 0.0094; bilateral DREADD rats: CLZ treatment, p_interaction_ = 0.012; CNO treatment, p_interaction_ = 0.037. Post hoc t-tests within each rat group: unilateral DREADD rats, SAL vs CLZ, p = 1.6×10^-9^; SAL vs CNO, p = 2.4×10^-8^; bilateral DREADD rats, SAL vs CLZ, p = 6.2×10^-6^; SAL vs CNO, p = 3.4×10^-5^; Fig. 2d).

Comparison of dHP fALFF between saline- and actuator-treated conditions using LME models (null hypothesis: β_1_ = 0, see Methods). T-tests for different treatment conditions in sham rats: SAL vs CLZ, p = 0.62, SAL vs CNO, p = 0.60. Two-way ANOVA with the virus type and treatment as main factors in DREADD rats: unilateral DREADD rats: CLZ treatment, p_interaction_ = 4.09×10^-4^; CNO treatment, p_interaction_ = 0.0094; bilateral DREADD rats: CLZ treatment, p_interaction_ = 0.012; CNO treatment, p_interaction_ = 0.037. Post hoc t-tests within each rat group is shown in **d)**: For unilateral DREADD rats, SAL vs CLZ, p = 1.6×10^- 9^; SAL vs CNO, p = 2.4×10^-8^. For bilateral DREADD rats, SAL vs CLZ, p = 6.2×10^-6^; SAL vs CNO, p = 3.4×10^-5^. n.s.: not significant; *: p<0.05; **: p<0.01; ***: p<0.001.

### dHP suppression induces brain-wide RSFC alterations

After confirming the suppression effects of DREADDs on local brain activities, we further assessed the impact of DREADDs on brain-wide connectivity. Consistent with the result of no significant fALFF suppression effect in the dHP in sham rats (Fig. 2d), we did not observe any significant RSFC change between saline- and either actuator-treated condition in sham rats (Fig. 3a, threshold: p < 0.05, FDR corrected), and the subthreshold off-target effects from the two actuators are similar (entry-to-entry Pearson correlation of their t matrices: r = 0.55, p = 1.9 × 10^-156^). To control for subthreshold off-target effects of CLZ and CNO, we created sham-derived RSFC difference matrices between saline and CLZ/CNO treated conditions. These maps were then respectively subtracted from both DREADD groups’ CLZ and CNO RSFC matrices. The resulting corrected DREADD RSFC values were compared against their corresponding saline RSFC values using LME models.

**Figure 3.**
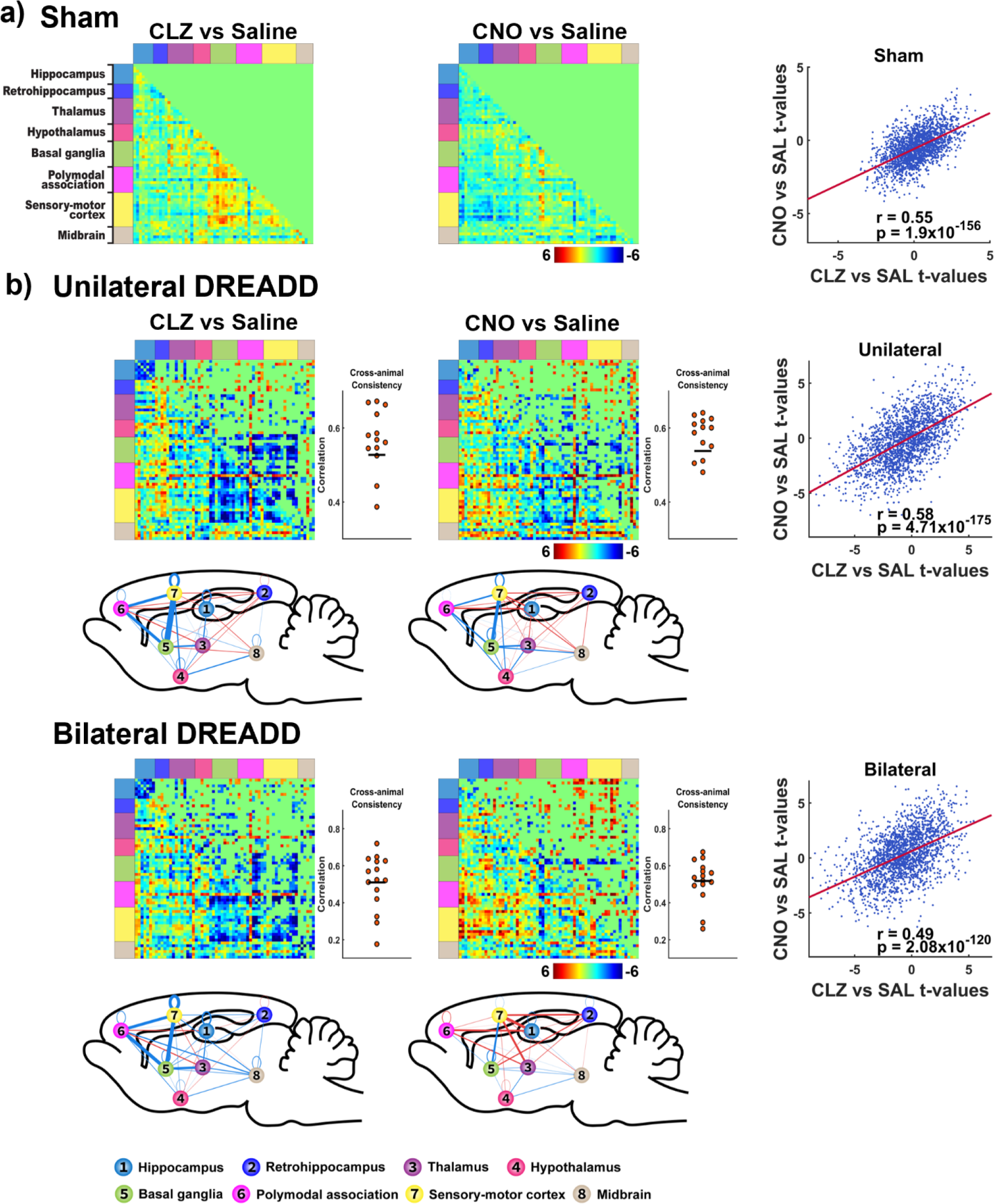
dHP suppression-induced alteration in brain-wide RSFC. a) RSFC differences between saline and actuator-treated conditions in sham rats. **b)** RSFC differences between saline- and actuator-treated conditions in each DREADD group after controlling for off-target actuator effects. For a) and b), lower triangles show non-thresholded t values without multiple comparison correction. Upper triangles show t values after FDR correction, p<0.05. Cross-animal consistency of DREADD-induced RSFC changes for each actuator is assessed by the entry-to-entry Pearson correlation between DREADD-induced RSFC difference matrix in each individual animal and the difference matrix averaged across all animals within the same group, shown on the right of the corresponding RSFC matrices (mean correlations cross animals: unilateral DREADD CLZ, r = 0.53; unilateral DREADD CNO, r = 0.54; bilateral DREADD CLZ, r = 0.51; bilateral DREADD CNO, r = 0.52). Brain system-level RSFC changes in different conditions are illustrated in the sagittal view of a glass brain slice beneath the corresponding RSFC-difference matrices. The thickness of the edge between numerically labeled nodes denotes the number of statistically significant RSFC changes identified between them. The color of the lines represents the nature of the overall RSFC changes: red indicates net positive RSFC changes, while blue indicates net negative RSFC change. Right scatter plots show entry-to-entry Pearson correlations of the t-value matrices between CLZ and CNO sessions. Sham rats: CLZ vs CNO, r = 0.55, p = 1.9 × 10^-156^. Unilateral DREADD rats: CLZ vs CNO, r = 0.58, 4.71 × 10^-175^. Bilateral DREADD rats: CLZ vs CNO, r = 0.49, p = 2.08 × 10^-120^.

Both CLZ and CNO elicited widely spread RSFC changes in both unilateral and bilateral DREADD groups (Fig. 3b). To assess cross-animal consistency of DREADD-induced RSFC changes, for each actuator we calculated the entry-to-entry Pearson correlation between the DREADD-induced RSFC difference matrix for each individual animal and the RSFC difference matrix averaged across all animals within the same group. In addition, given that both CLZ and CNO can be used to activate DREADDs, but through different pharmacokinetic mechanisms (Gomez et al., 2017), in DREADD animals, we first examined the potential difference in the effects on brain-wide RSFC between the two actuators. Our data show that within each DREADD group, the overall effects of the two actuators on RSFC matrices are consistent, reflected by significant entry-to-entry Pearson correlations of their corresponding t-value matrices (unilateral: r = 0.58, p = 4.71 × 10^-175^; bilateral: r = 0.49, p = 2.08 × 10^-120^) (Fig. 3b). This conclusion was replicated even when off-target effects of CLZ and CNO were not removed from DREADD groups (Fig. S3).

To gain a global view of the impact of DREADD suppression of dHP on RSFC, we divided the entire brain into eight systems: the hippocampus, retrohippocampus, thalamus, hypothalamus, basal ganglia, polymodal association cortex, sensory-motor cortex and midbrain. We then assessed RSFC changes within and between these systems. Our data revealed several prominent patterns of RSFC changes that are consistent between the unilateral and bilateral DREADD groups. First, DREADD suppression of dHP dampened the RSFC within the hippocampus system in both unilateral and bilateral DREADD groups, and this effect was more pronounced in the CLZ-treated condition (Fig. 3b). Second, both actuators strongly suppressed RSFC between the basal ganglia and cortex in both DREADD groups (Fig. 3b). Lastly, DREADD suppression of dHP also reduced the thalamic connectivity with basal ganglia (Fig. 3b). Taken together, our data indicate that dHP suppression produces a global impact on an extended hippocampal network comprised of the hippocampus, basal ganglia, thalamus and cortex. To further validate these results, we ran a group ICA on the rsfMRI data in sham rats. Two ICA components display clear involvement of hippocampus, basal ganglia, thalamus and cortex (Fig. S4), suggesting these brain areas indeed belong to an extended brain network.

### dHP suppression impairs spatial memory in animals

We next examined the effect of suppressing dHP activity on the animal’s memory performance at the behavioral level. We first asked whether any of the actuators affects locomotor activity, because it could influence the exploration behavior in Y-maze tests. At the same doses as in the fMRI experiment (CLZ, 0.1 mg/kg; CNO, 1mg/kg), the locomotion of sham rats was not affected by any of the actuators (Fig. S5). Likewise, DREADD rats did not display any difference in home-cage locomotion between saline- and actuator-treated conditions (Fig. S5). These data demonstrate that neither actuator affects locomotion in rats regardless of the presence of the hM4Di receptor.

The spatial recognition memory of the rat was measured using a two-trial Y-maze task (depicted in Fig. 4a) by recording the choice of first entry and percentage of time spent in the Novel Arm during Trial 2. Animals with intact spatial memory would exhibit preference to the Noval Arm to the Other ARM (Conrad et al.,1996). Across all treatment conditions, all sham rats exhibited strong preference to the Novel Arm in all conditions, indicated by both measures. First, in the choice of the first arm entry, a significant bias towards the Novel Arm was observed, as indicated by the results of the Chi-square tests: for saline treatment, X²(1, n = 8) = 4.5, p = 0.034. Compared with the saline condition, actuator- treated sham rats did not show significant difference in the choice of the first arm entry: SAL vs CLZ, X²(1, n = 16) = 1.07, p = 0.30; SAL vs CNO, X²(1, n = 16) = 1.07, p = 0.30 (Fig. 4b, left). Second, sham rats with saline treatment spent significant more time in the Novel Arm compared to a chance level (i.e. 33.3% of total duration, a one-sample t-test, t(7) = 2.59, p = 0.036), and this measure remains similar in both actuator-treated conditions (paired t-tests; SAL vs CLZ, t(7) = 1.08, p = 0.30; SAL vs CNO, t(7) = 0.11, p = 0.92, Fig. 4c). Taken together, these results indicate that the spatial memory of sham animals remained intact when treated with actuators alone.

**Figure 4.**
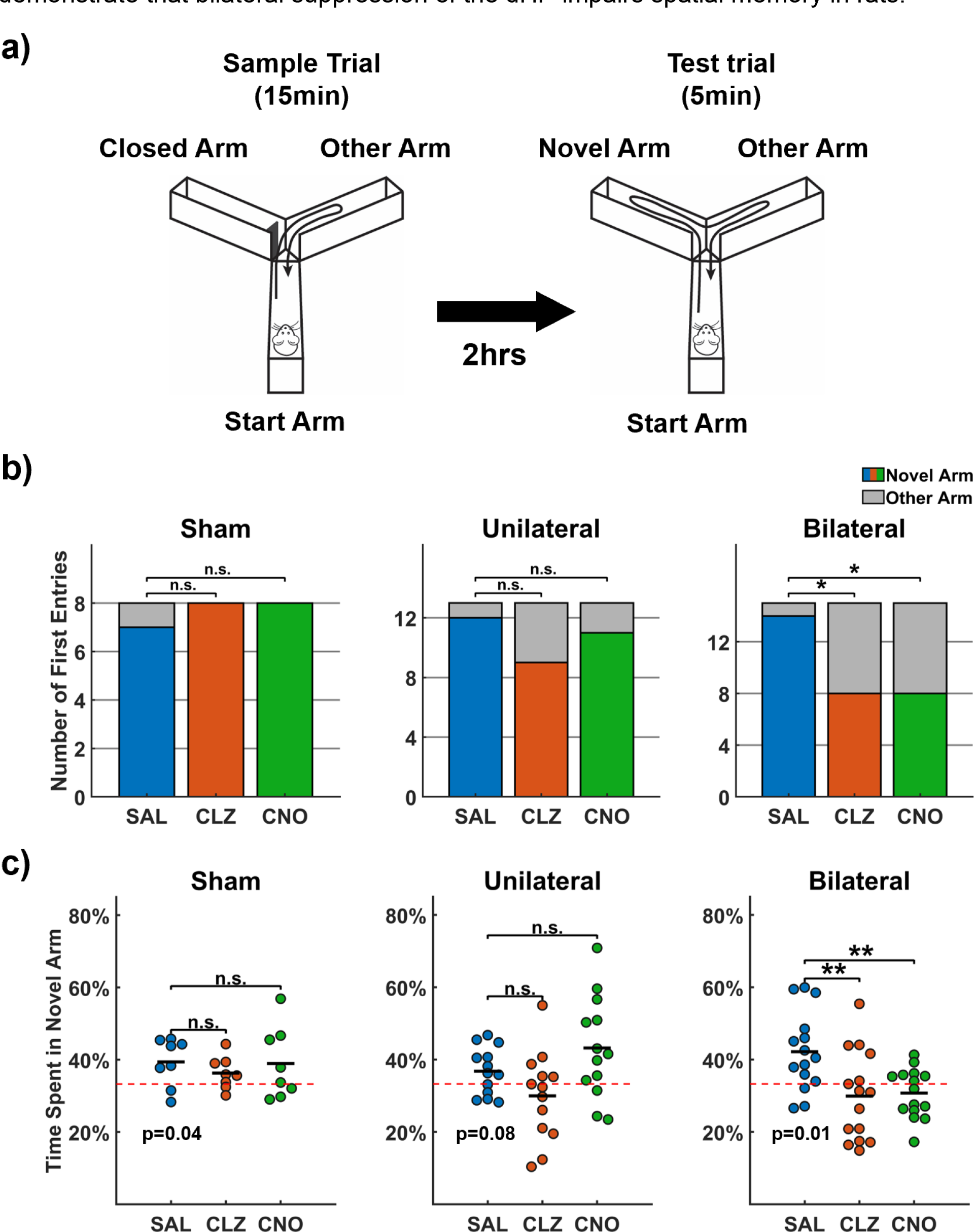
Effects of DREADD suppression on Y-maze tests. a) Diagram illustration of the Y maze test. **b)** Analysis of first-arm entry in Y maze test. The choice of first-arm entry is color coded (Novel Arm: colored; Other Arm: grey). For saline-treated conditions, the number of rats entered the Novel Arm first was compared with that of a random chance by Chi-square test: sham rats: X^2^(1, n = 8) = 4.5, p = 0.034; unilateral DREADD rats: X^2^(1, n = 13) = 9.3, p = 0.0023; bilateral DREADD rats: X^2^(1, n = 15) = 11.3, p = 7.9 × 10^-4^. For actuator-treated conditions, the relationship between treatment (saline vs. actuator) and the number of rats entered the Novel Arm first was examined by 2-by-2 chi-square test. For sham rats: SAL vs CLZ, X^2^(1, n = 16) = 1.07, p = 0.30; SAL vs CNO, X^2^(1, n = 16) = 1.07, p = 0.30. For unilateral DREADD rats: SAL vs CLZ, X^2^(1, n = 26) = 2.23, p = 0.14; SAL vs CNO, X^2^(1, n = 26) = 0.38, p = 0.54. For bilateral DREADD rats: SAL vs CLZ, X^2^(1, n = 30) = 6.14, p = 0.013; SAL vs CNO, X^2^(1, n = 30) = 6.14, p = 0.013. **c)** Analysis of percentage of time spent in the Novel Arm. The red dashed line at 33.3% indicates the time spent in the Novel Arm by chance (3 arms in total). For saline-treated conditions, data was compared with 33.3% by one-sample t-test. Sham rats, p = 0.036; unilateral DREADD rats, p = 0.078; bilateral DREADD rats, p = 0.0069. In each rat group, data of actuator-treated conditions were compared with data of saline-treated condition through paired t-tests. Sham rats: SAL vs CLZ, p = 0.41; SAL vs CNO, p = 0.90. Unilateral DREADD rats: SAL vs CLZ, p = 0.13; SAL vs CNO, p = 0.16. Bilateral DREADD rats: SAL vs CLZ, p = 0.0021; SAL vs CNO, p = 0.0039. n.s.: not significant; *: p< 0.05; **: p<0.01.

For DREADD rats treated with saline, the spatial memory also remained intact (first entry: Chi-square test, unilateral: X^2^(1, n = 13) = 9.3, p = 0.0023; bilateral: X^2^(1, n = 15) = 11.3, p = 7.9 × 10^-4^; time spent in the Novel Arm: one sample t-tests, unilateral: t(12) = 1.93, p = 0.079; bilateral: t(14) = 3.16, p = 0.0069). For unilateral DREADD rats, after CLZ or CNO injection, the distribution of the first arm entry was not significantly different from that of saline injection (2-by-2 chi-square test: SAL vs CLZ, X^2^(1, n = 26) = 2.23, p = 0.14; SAL vs CNO, X^2^(1, n = 26) = 0.38, p = 0.54). Consistently, the time spent in the Novel Arm after actuator injection was also indifferent from that after saline injection (paired t- tests: SAL vs CLZ, p = 0.13; SAL vs CNO, p = 0.16) (Figs. 4b-c, Fig. S6), indicating DREADD-induced suppression of dHP in one hemisphere is not sufficient to cause significant spatial-memory loss. In contrast, for bilateral DREADD rats, both actuators caused significant loss of memory in the Y-maze test, as evidenced by the lack of preference for the Novel arm during their first entry: 8 rats entered the Novel Arm, and 7 rats entered the Other Arm under both CLZ and CNO treatments. This distribution significantly differed from that observed with saline treatment (2-by-2 chi-square test: SAL vs CLZ, X^2^(1, n = 30) = 6.14, p = 0.013; SAL vs CNO, X^2^(1, n = 30) = 6.14, p = 0.013).

Additionally, both actuators caused significant reduction of the time spent in the Novel Arm relative to the saline treatment (paired t-tests: CLZ vs SAL, t(14) = -2.89, p = 0.007; CNO vs SAL, t(14) = -3.46, p = 0.002) (Figs. 4b-c, Fig. S6). These results collectively demonstrate that bilateral suppression of the dHP impairs spatial memory in rats.

Next, we tested the hypothesis that changes in memory-related behavior are related to RSFC changes in brain networks. To do this, for each treatment condition RSFC matrices from repeated scans were first averaged for each animal. Then, for each actuator we calculated the correlation between actuator-induced RSFC changes for each connection in the whole-brain network and changes in time spent in the Novel Arm across all animals within the same group. This correlation analysis was conducted across all three groups of animals, comparing how each connection’s RSFC change corresponded to the behavioral change in response to each actuator treatment. Fig. 5 shows behavior-RSFC correlation matrices in CLZ- and CNO-treated conditions in all three groups of animals. In sham rats, these behavior-RSFC correlations derived from CLZ- and CNO-treated conditions are weakly anti-correlated (r = -0.13, p = 6.19 ×10^-9^) (Fig. 5, upper panel). For unilateral DREADD rats, the two behavior-RSFC correlation matrices from CLZ- and CNO-treated conditions are uncorrelated (r = 0.02, p = 0.46) (Fig. 5, middle panel). In contrast, the behavior-RSFC correlation was highly correlated in CLZ- and CNO-treated conditions for the bilateral DREADD group (r = 0.65, p = 1.38 ×10^-235^) (Fig. 5, lower panel), indicating CLZ- and CNO-generated memory deficits share a similar underlying brain connectivity change architecture.

**Figure 5.**
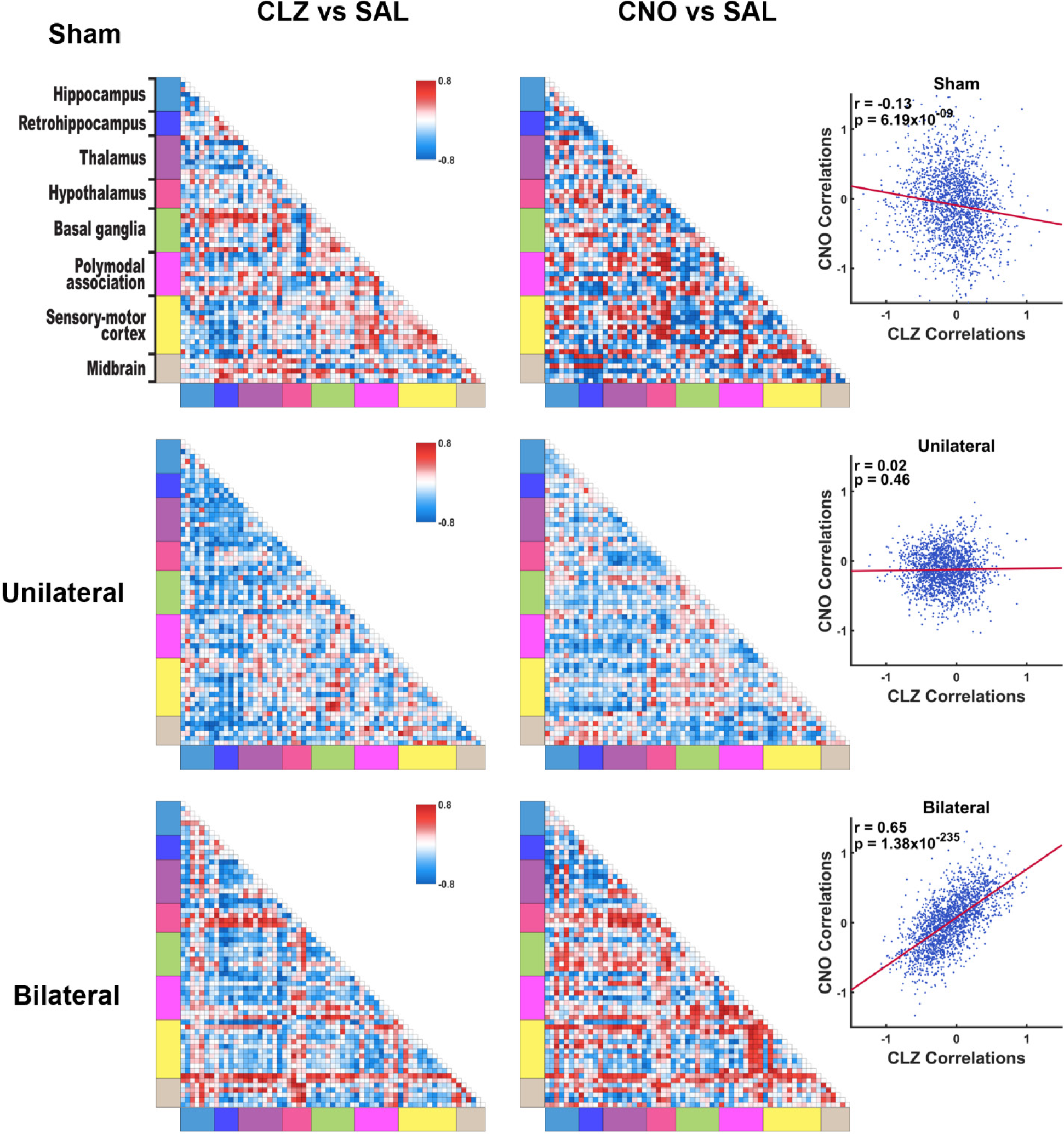
Behavior-RSFC correlations. Actuator-induced changes in percentage of time spent in the Novel Arm in Y-maze tests were correlated to the same actuator-induced RSFC changes for individual connections across animals within the same group. Matrices on the left show these correlations for all connections in the whole-brain network. Scatter plots on the right show the correlation of the z-transformed behavior-RSFC correlation matrices between CLZ- and CNO-treated conditions. Sham rats: r = -0.13, p = 6.19 x10^-9^. Unilateral DREADD rats: r = 0.02, p = 0.46. Bilateral DREADD rats: r = 0.65, p = 1.38 x 10^-235^.

### Brain network-level interpretation of DREADD-induced impaired memory

Considering the robust behavioral effect (Fig. 4) and consistent behavior-RSFC correlations between the two actuators in bilateral DREADD animals, we further explored the network-level interpretation of impaired memory in this group. For this purpose, we constructed the brain network associated with the Y-maze memory performance by selecting connections exhibiting significant RSFC-behavior correlations under both CLZ and CNO-treated conditions (p<0.05 for each condition). This thresholding process generated a network comprised of 45 connections. Given that all of these connections showed consistent trends in RSFC-behavior correlations between the two actuator- treated conditions in the bilateral DREADD group (Fig. S7), we calculated one aggregated correlation coefficient between RSFC changes and behavior change (time spent in the novel arm) by pooling these data from both CNO- and CLZ-treated conditions for all individual 45 connections, as shown in Fig. 6. In this network model, RSFC changes between the sensory-motor cortex (predominantly confined to the visual cortex (VIS)) and hippocampus, as well as thalamus, positively correlate with the memory performance change in bilateral DREADD rats, indicating that decreased RSFC is associated with poorer memory performance. RSFC changes between the thalamus and hypothalamus, including anterior dorsal/ventral thalamus nuclei (AV and AD) and the mammillary complex (MAM), also positively correlate with memory performance changes. In addition, RSFC changes within the midbrain show positive correlations with the memory change. Conversely, multiple thalamic nuclei display negative correlation with memory performance through their RSFC with the retrohippocampus, polymodal association cortex, basal ganglia and sensory-motor cortex (i.e. stronger RSFC is associated with worse memory performance). Taken together, our data provide a comprehensive brain network model that is linked to memory performance alteration, and support the notion that the memory function involves a properly coordinated network spanning the entire brain.

**Figure 6.**
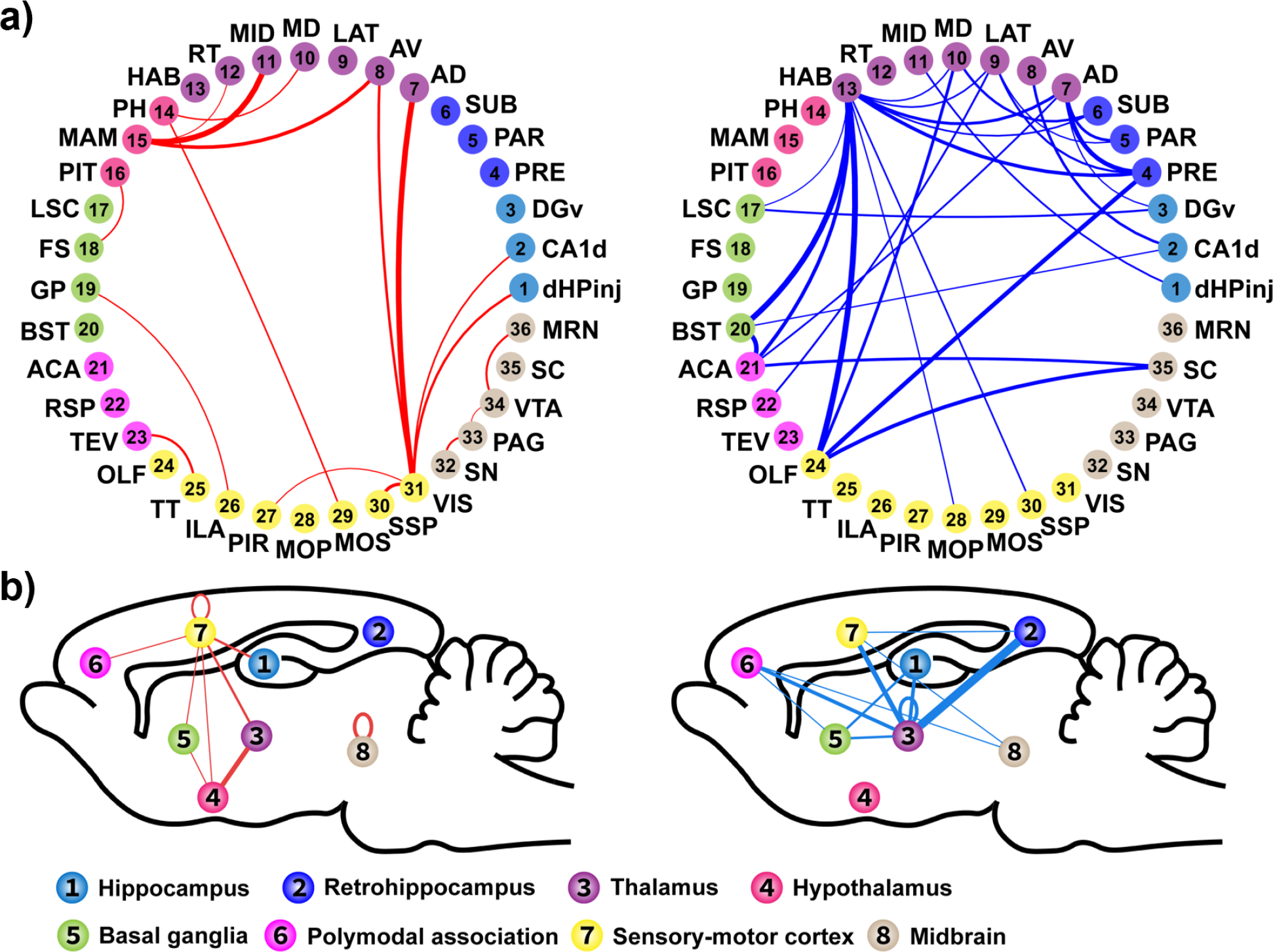
A brain network model to interpret DREADD-induced memory performance change. a) Memory-related RSFC organized by constituent brain regions. Connections showing significant behavior-RSFC correlations in both CLZ- and CNO-treated conditions (p<0.05 for each condition, joint probability < 0.0025) in the bilateral DREADD group are plotted in the circle graph. Circular dots and curved lines represent brain regions and the connections between them, respectively. The line thickness reflects the strength of behavior-RSFC correlation, which was calculated by pooling data from both actuator-treated conditions. Red and blue lines indicate positive and negative correlations, respectively. Abbreviations: dHPinj, dHP injection site; CA1d, CA1 dorsal; DGv, dentate gyrus ventral; PRE, presubiculum; PAR, parasubiculum; SUB, subiculum; AD, anterior dorsal thalamic nucleus; AV, anterior ventral thalamic nucleus; LAT, lateral thalamic area; MD, medial dorsal thalamus; MID, midline thalamic nuclei; RT, reticular thalamic nucleus; HAB, habenula; PH, posterior hypothalamic nucleus; MAM, mammillary complex; PIT, pituitary; LSC, lateral septum complex; FS, fundus of the striatum; GP, globus pallidus; BST, stria terminalis; ACA, anterior cingulate cortex; RSP, retrosplenium complex; TEV, ventral temporal association areas; OLF, olfactory bulb; TT, taenia tecta; ILA, infralimbic area; PIR, piriform area; MOP, primary motor cortex; MOS, secondary motor cortex; SSP, primary somatosensory cortex; VIS, visual cortex; SN, substantia nigra; PAG, periaqueductal gray; VTA, ventral tegmental area; SC, superior colliculus; MRN, mesencephalic reticular nucleus. **b)** Memory-related RSFC organized by conglomerated brain systems. RSFC between brain regions shown in **a)** were grouped and summed according to the brain systems they belong to. The line thickness indicates the number of connections between the two systems that are involved in memory loss, and positive (upper panel) and negative (lower panel) behavior-RSFC correlations are plotted separately in the sagittal view of the glass brain.

## Discussion

Studying the brain-behavior relationship is a central theme in neuroscience. Conventional methods use focal stimulation (e.g. electrical stimulation) or lesion to investigate how a brain region is involved in certain behavior. Although situationally useful, these historical approaches are notoriously unspecific. More modern neuroscience approaches such as optogenetics and chemogenetics allow for more spatially precise brain manipulations with cell-type specificity, which enable scientists to intentionally explore the role of specific neural circuits in behavior. However, these approaches alone are not sufficient for the investigation of large-scale brain networks, in which coordinated activity is key to mediate complex behaviors (Yamashita et al., 2015; Liégeois et al., 2019). To tackle this issue, here we incorporated chemogenetic manipulation, rsfMRI and behavioral tests to study how changes in memory performance are caused by targeted perturbation of an extended hippocampal network in a rodent model. Our data reveal that suppressing dHP activity has a significant impact on local and global connectivity in a large-scale brain network, along with the memory performance. Interestingly, neural connections associated with the memory behavioral changes are not confined to the hippocampal formation itself but are widely distributed across other, more distant brain systems such as the cortex, thalamus, and hypothalamus. This result suggests that the impact of dHP inhibition on memory behavior is mediated by complex interactions between multiple brain systems. Additionally, we showed that two different DREADD actuators, CLZ and CNO, induced consistent effects on both brain networks and behavior tests, despite having distinct metabolic processes in the body (Gomez et al., 2017; Manvich et al., 2018) and different off-target binding profiles in the brain (Jendryka et al., 2019). These data highlight the utility of the DREADD-fMRI technology and its application in studying brain-behavior relationship.

### Relating brain network RSFC to spatial memory-related behavior

The brain network possesses small-world properties: some brain regions have more extensive connections and play a central role in neuronal communication and integration (Oldham & Fornito, 2019). Thus, inhibiting the dHP, a central node in the network for memory processing (Geib et al., 2017), is expected to have an overall impact on brain-wide networks and memory-related behavior. Our findings are consistent with this theoretical expectation, as we observed significant network alterations not only within the hippocampal formation but also in numerous areas essential for learning and memory (Fig. 3b). As a result, these network/circuit-level changes can contribute to altered performances in the spatial memory test in animals.

To further investigate this issue, we first hypothesize that DREADD-induced connectivity change between the dHP and visual cortex influences the spatial memory performance. Anatomically, the dHP receives inputs from all sensory cortices via the EC and subsequently processes the information in neural ensembles through the trisynaptic pathway of EC-DG-CA3-CA1 (Amaral et al., 2007). During visual field mapping, the hippocampus establishes a spatial representation for the visual cortex (Silson et al., 2021; Hindy et al., 2016), and the interplay between the hippocampus and visual cortex facilitates memory consolidation during slow-wave sleep (Rothschild et al., 2017; Yu et al., 2023). In addition, it has been shown that impaired hippocampal-visual cortex connectivity is related to compromised spatial representation precision in hippocampus (Domenico et al., 2021). In the current study, the long-lasting effects of actuators can compromise the memory consolidation process during the two-hour ITI in the Y-maze test. Consistent with this hypothesis, our data show that deactivation of the dHP markedly influences its RSFC with the visual cortex, and this RSFC change is significantly associated with changes in spatial memory performance (Fig. 6).

We also hypothesize that the RSFC between nodes in the Papez circuit is essential for spatial memory performance. The Papez circuit is a critical brain network involved in memory process, comprising the hippocampal formation, mammillary bodies (MAM), anterior thalamic nuclei (ATN), and the retrosplenium (RSP). Anatomically, the ATN receives its main ascending projection from the subcortical MAM, which in turn receives input from the hippocampal formation. The ATN sends efferent projections to the RSP, which projects back to the hippocampal formation, completing the Papez circuit (Jones & Witter, 2007; Aggleton et al., 2016). Our data support this hypothesis, showing that changes in RSFC involving the MAM and anterior dorsal/ventral thalamic nuclei (AD and AV) are significantly associated with altered spatial memory performance. These results are well in line with previous literature findings. The involvement of ATN in spatial memory has been well documented (Perry & Mitchell, 2019). Lesions to the ATN are associated with impair performance in spatial memory in rodents (Lopez et al., 2009). A recent human fMRI study also showed that the ATN is activated during visual information encoding (Sweeney-Reed et al., 2021). Besides, we found that changes in spatial memory performance are positively correlated with RSFC changes between the visual cortex and the AD and AV. Although the visual cortex is not considered part of the Papez circuit, this correlation might be mediated by other nodes within the memory-related network.

Moreover, we identified the MID as a participating node in the memory-related network, consistent with the literature that the nucleus reuniens (RE), a part of MID, receives input from the MAM and serves as a relaying center for memory information (McKenna & Vertes, 2004; Mathiasen et al., 2020). Other nodes found to be involved in the spatial memory-associated network such as the ventral tegmental area (VTA) are also in agreement with the literature report (Vann, 2009).

In addition to testing specific hypotheses, the whole-brain coverage provided by fMRI makes the DREADD-fMRI-behavior approach a valuable tool for identifying circuits that may have previously been overlooked. In the current study we discovered that spatial memory performance is positively associated with RSFC between visual and odor processing regions. Specifically, this includes previously unreported connections between the VIS and the piriform area (PIR), as well as the ventral temporal association areas (TEV) and taenia tecta (TT). The VIS and TEV are critical for visual processing (Kravitz et al., 2013), while the TT and PIR are involved in processing olfactory information (Wilson & Stevenson, 2003). Efficient communication between these sensory systems likely creates a more comprehensive representation of the environment, facilitating better encoding and retrieval of spatial information. In our experiment, although the Y-maze apparatus was cleaned with alcohol after each use, some residual odors may have remained detectable to the rats. These findings suggest that the integration of sensory information might enhance spatial memory.

Collectively, our results present an extensive memory-associated network that extends well beyond the hippocampal formation itself and underscores the importance of coordinated activity within this network in memory process. This finding highlights the power of whole-brain analysis in studying the neural mechanisms underlying complex behaviors. This advantage has also been demonstrated in previous studies. For instance, (Tu et al., 2021) combined fMRI and chemogenetics with behavioral tests and found that RSFC within the default mode network (DMN) is linked to restfulness behavior, illustrating how the whole-brain approach can uncover critical functional connections for understanding behavior. Similarly, Yamashita et al. (2015) highlighted the role of the frontal-parietal network in task execution, further emphasizing the efficacy of whole-brain analyses in identifying networks involved in cognitive processes. These findings showcase the potential of combining whole-brain imaging and neuromodulation tools to advance our understanding of brain function by revealing how circuits can causally contribute to behavior and task performance. However, it is important to note that not all connections exhibiting RSFC-behavior correlations in our study play a causal role in memory performance, as RSFC changes in some connections can covary with behavior due to indirect effects from other critical nodes.

### DREADDs and actuator effects

In the current study, we utilized the DREADD technique to suppress brain activity by taking advantage of its reversibility and long-lasting effect. Nonetheless, the off-target effect of actuators used in DREADDs may represent a limitation of this system. CLZ is a sedative drug that treats schizophrenia by binding to several endogenous targets, including serotoninergic, dopaminergic and adrenergic receptors (Martinez et al., 2019). Similarly, CNO binds to a wide range of receptors, such as adrenergic, histaminergic and muscarinic receptors, with relatively high affinity (Jendryka et al., 2019). To minimize the potential off-target effects, we administered low-dose CNO and CLZ (CNO, 1 mg/kg; CLZ, 0.1 mg/kg), as recommended by literature studies (Jendryka et al., 2019; Gomez et al., 2017). In addition, our results were cross validated using both actuators (CLZ and CNO). For both actuators, we observed consistent subthreshold alterations in brain-wide RSFC in sham rats (Fig. 3a). In addition, within each DREADD group, both actuators had similar impact on RSFC (Fig. 3b). In the bilateral DREADD group, both actuators impaired the spatial memory of rats in a similar manner (Fig. 4). Moreover, our data show that significant correlations between behavioral and RSFC changes in both actuator-treated conditions exhibit similar trends, indicating that CLZ and CNO induce comparable effects on the brain-behavior relation (Fig. S7). Recently, Nagai et al. (2020) introduced deschloroclozapine (DCZ) as another potent DREADD actuator. DCZ is noted for its stability and reduced off-target effects. However, its pharmacokinetics indicate that the DCZ concentrations in the plasma and brain decrease significantly within two hours post-injection, and it is metabolized into C21 and DCZ-N-oxide. Some studies have reported equivalent or superior results with DCZ compared to other actuators (Ferrari et al., 2022; Nentwig et al., 2022; Shimizu et al., 2023). Therefore, DCZ could be a useful alternative worth considering in future research.

## Summary

By combining DREADDs, fMRI, and behavioral tests, we have demonstrated that suppressing a central node in the memory-related network induces whole-brain RSFC changes, and a subset of these changes correlate with relevant behavioral changes. Based on the associated changes triggered by the same experimental manipulation, we constructed a brain network model that is linked to memory performance. These results provide a broader and more comprehensive framework for better understanding of the neural mechanism of memory function, particularly from a network perspective.

## Acknowledgments

The present study was partially supported by National Institute of Neurological Disorders and Stroke (R01NS085200). The content is solely the responsibility of the authors and does not necessarily represent the official views of the National Institutes of Health.

