## Supplemental Document for "Exploring memory-related network via dorsal hippocampus suppression"

**Acknowledgments**

The present study was partially supported by National Institute of Neurological Disorders and Stroke (R01NS085200). The content is solely the responsibility of the authors and does not necessarily represent the official views of the National Institutes of Health.

### Supplementary figures

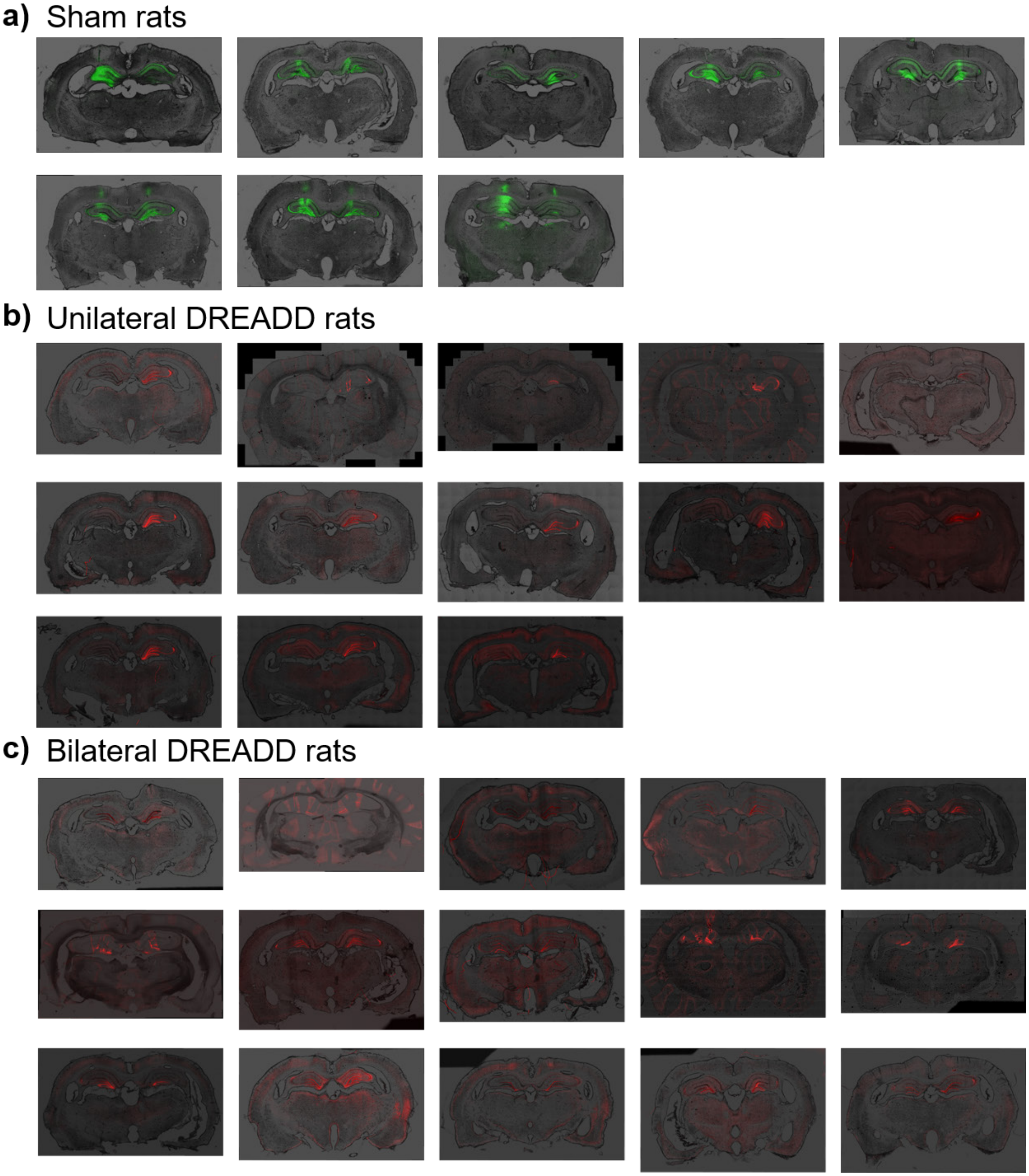


**Figure S1. Histology of all animals a)** Sham rats. **b)** Unilateral DREADD rats**. c)** Bilateral DREADD rats.

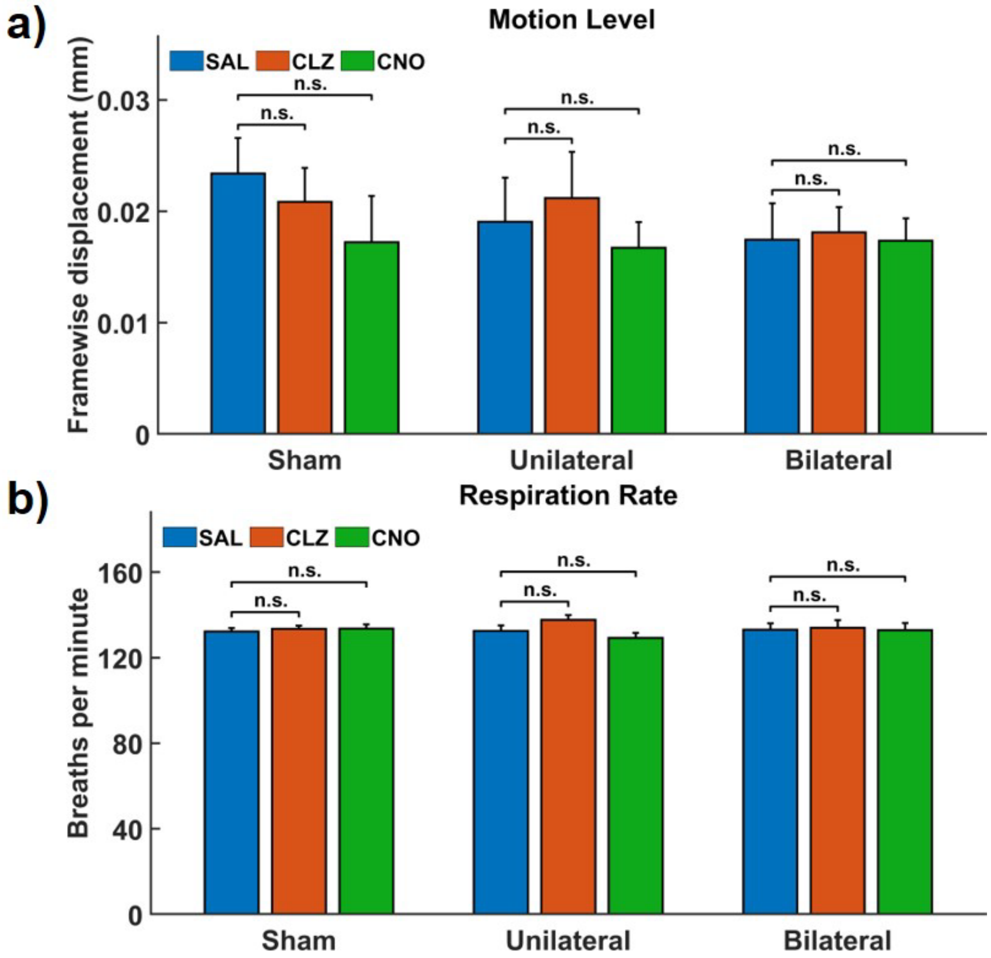


**Figure S2. Motion level and respiration rate during scanning. a)** Motion level. T-tests for different conditions in sham rats show no difference in motion level (SAL vs CLZ, $p = 0.16$, SAL vs CNO, $p = 0.13$. For DREADD rats, the interaction terms of two-way ANOVAs with virus type and treatment as factors were analyzed. Unilateral DREADD rats: SAL vs CLZ, $p_{interaction} = 0.37$, SAL vs CNO, $p_{interaction} = 0.53$. Bilateral DREADD rats: SAL vs CLZ, $p_{interaction} = 0.46$, SAL vs CNO, $p_{interaction} = 0.14$. **b)** Respiration rate. T-tests for different conditions in sham rat show SAL vs CLZ, $p = 0.60$, SAL vs CNO, $p = 0.61$. For DREADD rats, the interaction terms of two-way ANOVAs with virus type and treatment as factors were analyzed. Unilateral DREADD rats: SAL vs CLZ, $p_{interaction} = 0.42$, SAL vs CNO, $p_{interaction} = 0.25$. Bilateral DREADD rats: SAL vs CLZ, $p_{interaction} = 0.72$, SAL vs CNO, $p_{interaction} = 0.84$. n.s. not significant.

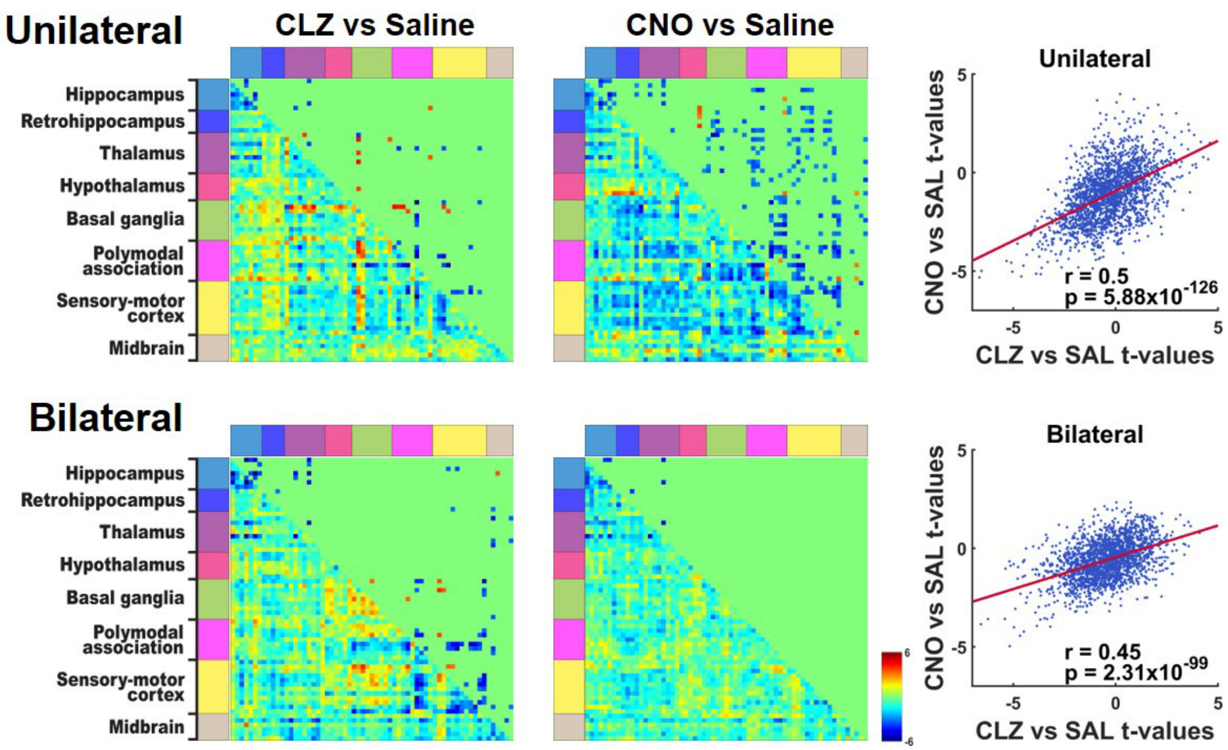


**Figure S3. dHP suppression-induced alteration in brain-wide RSFC (without subtracting 'sham differences').** T-tests of RSFC between saline- and actuator-treated conditions in DREADD groups. In this analysis, RSFC differences between the actuator- and saline-treated conditions in the sham group were not subtracted out from the DREADD data. Lower triangles show non-thresholded t values. Upper triangles show t values thresholded at $p < 0.05$, after FDR correction. Right scatter plots show correlations of the t-value matrices between CLZ and CNO administered sessions within the same DREADD groups. Unilateral DREADD rats: CLZ vs CNO, $r = 0.5$, $5.88 \times 10^{-126}$. Bilateral DREADD rats: CLZ vs CNO, $r = 0.45$, $p = 2.31 \times 10^{-99}$.

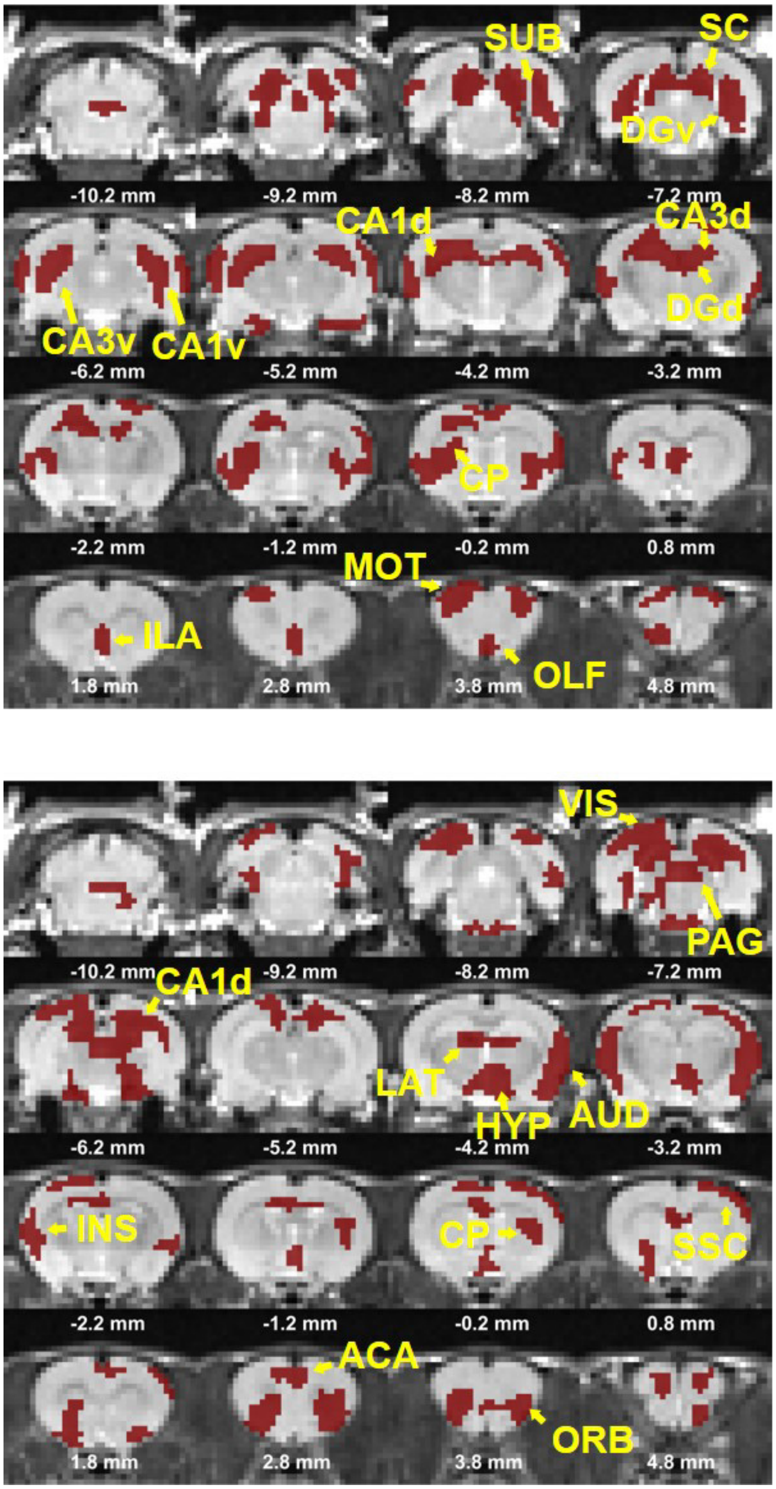


**Figure S4. ICA components of the hippocampus network.** ICA was performed on the rsfMRI data collected from the sham group, generating 20 independent components. Two components with the hippocampal network are displayed.

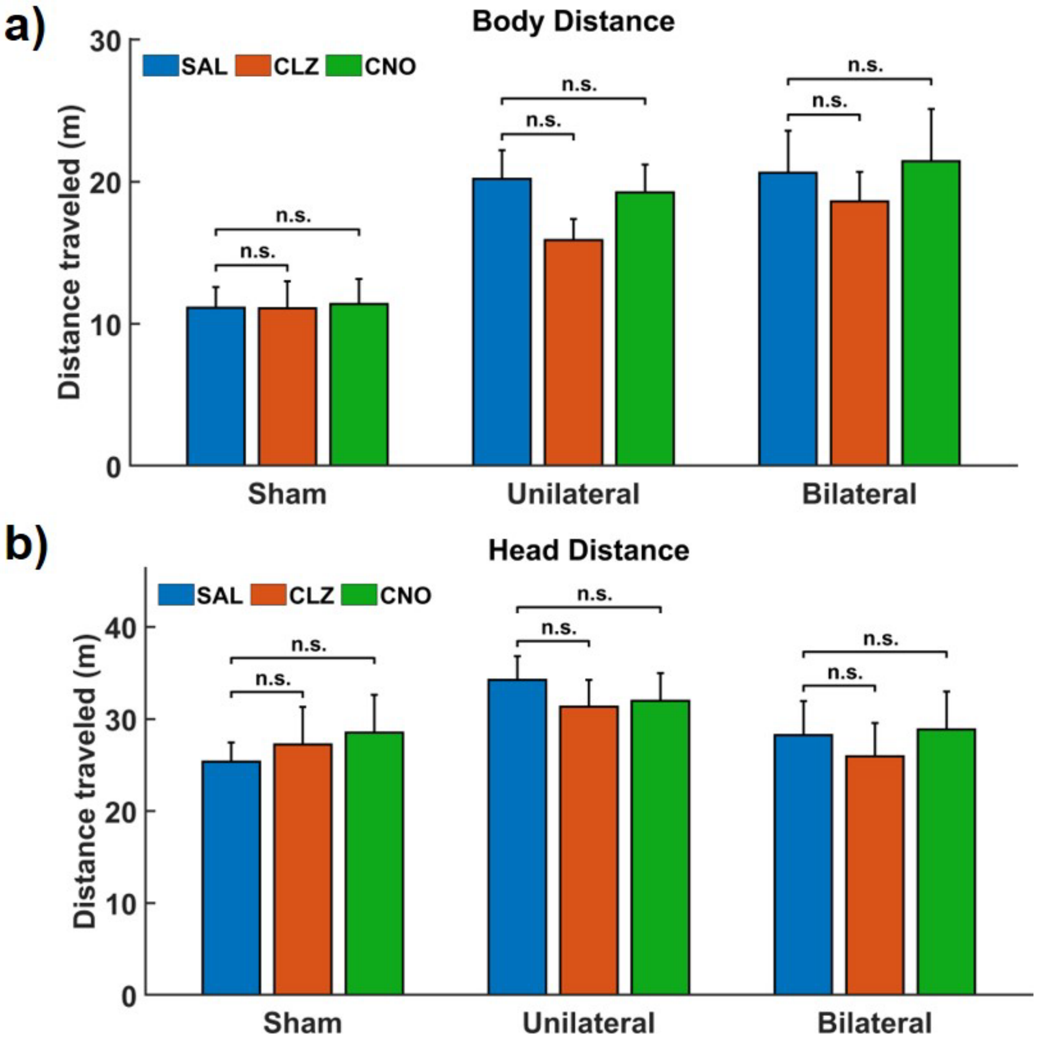


**Figure S5. Home cage locomotion after administration of saline or actuators. a)** Total body distance. T-tests for different conditions in sham rats: SAL vs CLZ, $p = 0.99$; SAL vs CNO, $p = 0.84$. For DREADD rats, the interaction terms of two-way ANOVAs with virus type and treatment as factors were analyzed. Unilateral DREADD rats: SAL vs CLZ, $p_{interaction} = 0.23$; SAL vs CNO, $p_{interaction} = 0.67$. Bilateral DREADD rats: SAL vs CLZ, $p_{interaction} = 0.45$; SAL vs CNO, $p_{interaction} = 0.87$. **b)** Total head distance. T-tests for different conditions in sham rats: SAL vs CLZ, $p = 0.69$; SAL vs CNO, $p = 0.49$. For DREADD rats, the interaction terms of two-way ANOVAs with virus type and treatment as factors were analyzed. Unilateral DREADD rats: SAL vs CLZ, $p_{interaction} = 0.45$; SAL vs CNO, $p_{interaction} = 0.35$. Bilateral DREADD rats: SAL vs CLZ, $p_{interaction} = 0.42$; SAL vs CNO, $p_{interaction} = 0.58$. n.s., not significant.

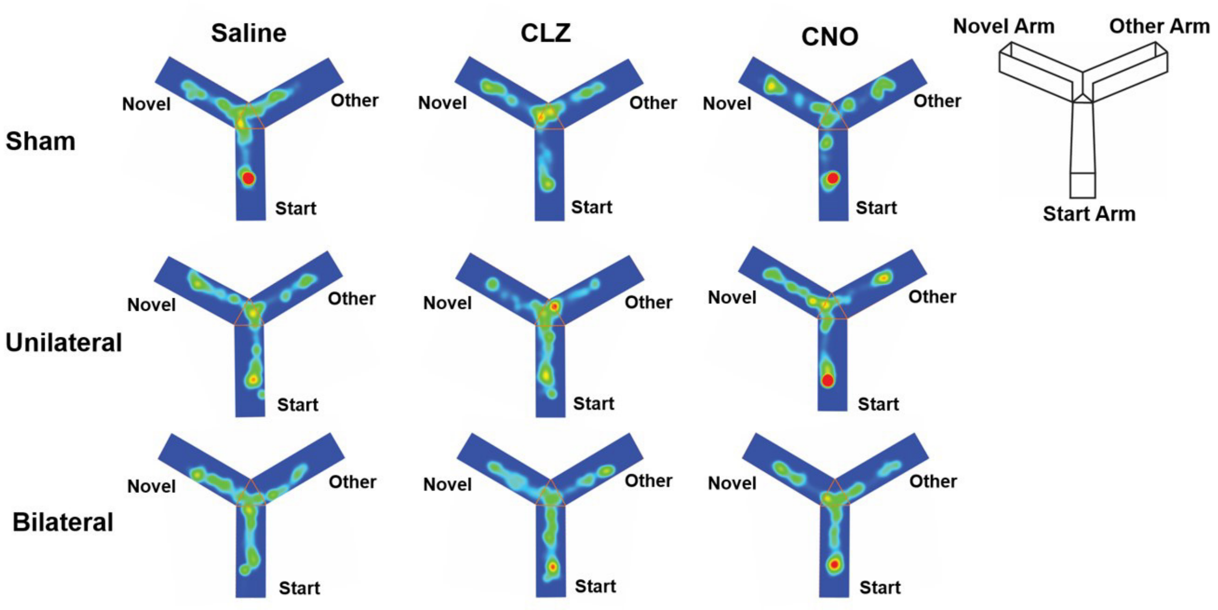


**Figure S6. Representative heatmaps in the second trial of the Y-maze test.** The representative heatmaps were picked from the trials showing the percentage of time spent in the Novel Arm that was the closest to the averaged value in same condition of the same group.

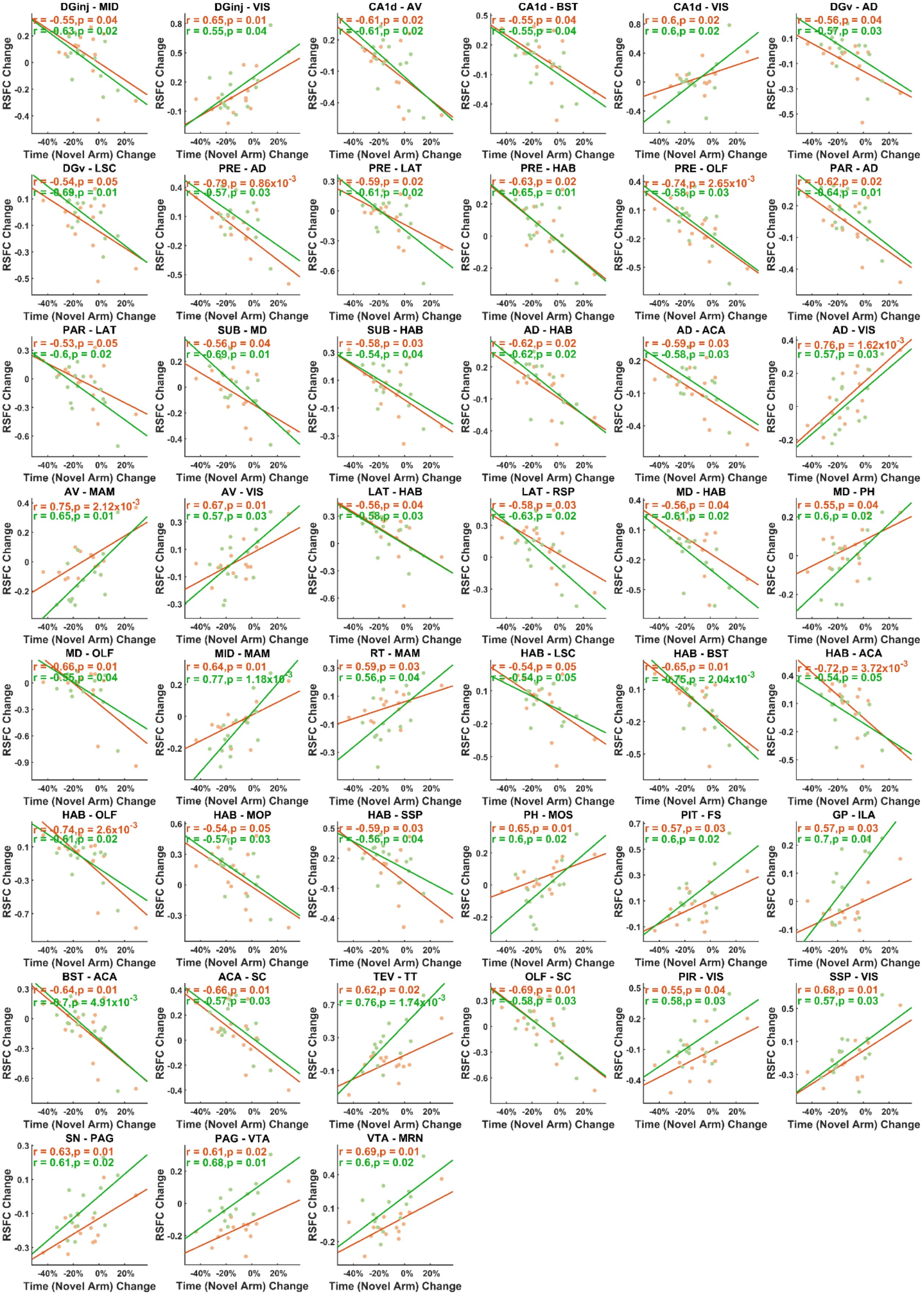

DGinj - MID
r = -0.55,p = 0.04
r = -0.63,p = 0.02
DGinj - VIS
r = 0.65,p = 0.01
r = 0.55,p = 0.04
CA1d - AV
r = -0.61,p = 0.02
r = -0.61,p = 0.02
CA1d - BST
r = -0.55,p = 0.04
r = -0.55,p = 0.04
CA1d - VIS
r = 0.6,p = 0.02
r = 0.6,p = 0.02
DGv - AD
r = -0.56,p = 0.04
r = -0.57,p = 0.03
DGv - LSC
r = -0.54,p = 0.05
r = -0.69,p = 0.01
PRE - AD
r = -0.79,p = 0.86x10-3
r = -0.57,p = 0.03
PRE - LAT
r = -0.59,p = 0.02
r = -0.61,p = 0.02
PRE - HAB
r = -0.63,p = 0.02
r = -0.65,p = 0.01
PRE - OLF
r = -0.74,p = 2.65x10-3
r = -0.58,p = 0.03
PAR - AD
r = -0.62,p = 0.02
r = -0.64,p = 0.01
PAR - LAT
r = -0.53,p = 0.05
r = -0.6,p = 0.02
SUB - MD
r = -0.56,p = 0.04
r = -0.69,p = 0.01
SUB - HAB
r = -0.58,p = 0.03
r = -0.54,p = 0.04
AD - HAB
r = -0.62,p = 0.02
r = -0.62,p = 0.02
AD - ACA
r = -0.59,p = 0.03
r = -0.58,p = 0.03
AD - VIS
r = 0.76,p = 1.62x10-3
r = 0.57,p = 0.03
AV - MAM
r = 0.75,p = 2.12x10-3
r = 0.65,p = 0.01
AV - VIS
r = 0.67,p = 0.01
r = 0.57,p = 0.03
LAT - HAB
r = -0.56,p = 0.04
r = -0.58,p = 0.03
LAT - RSP
r = -0.58,p = 0.03
r = -0.63,p = 0.02
MD - HAB
r = -0.56,p = 0.04
r = -0.61,p = 0.02
MD - PH
r = 0.55,p = 0.04
r = 0.6,p = 0.02
MD - OLF
r = -0.66,p = 0.01
r = -0.55,p = 0.04
MID - MAM
r = 0.64,p = 0.01
r = 0.77,p = 1.18x10-3
RT - MAM
r = 0.59,p = 0.03
r = 0.56,p = 0.04
HAB - LSC
r = -0.54,p = 0.05
r = -0.54,p = 0.05
HAB - BST
r = -0.65,p = 0.01
r = -0.75,p = 2.04x10-3
HAB - ACA
r = -0.72,p = 3.72x10-3
r = -0.54,p = 0.05
HAB - OLF
r = -0.74,p = 2.6x10-3
r = -0.61,p = 0.02
HAB - MOP
r = -0.54,p = 0.05
r = -0.57,p = 0.03
HAB - SSP
r = -0.59,p = 0.03
r = -0.56,p = 0.04
PH - MOS
r = 0.65,p = 0.01
r = 0.6,p = 0.02
PIT - FS
r = 0.57,p = 0.03
r = 0.6,p = 0.02
GP - ILA
r = 0.57,p = 0.03
r = 0.7,p = 0.01
BST - ACA
r = -0.64,p = 0.01
r = -0.7,p = 4.91x10-3
ACA - SC
r = -0.66,p = 0.01
r = -0.57,p = 0.03
TEV - TT
r = 0.62,p = 0.02
r = 0.76,p = 1.74x10-3
OLF - SC
r = -0.69,p = 0.01
r = -0.58,p = 0.03
PIR - VIS
r = 0.55,p = 0.04
r = 0.58,p = 0.03
SSP - VIS
r = 0.68,p = 0.01
r = 0.57,p = 0.03
SN - PAG
r = 0.63,p = 0.01
r = 0.61,p = 0.02
PAG - VTA
r = 0.61,p = 0.02
r = 0.68,p = 0.01
VTA - MRN
r = 0.69,p = 0.01
r = 0.6,p = 0.02
RSFC Change
Time (Novel Arm) Change

**Figure S7**. **Behavior-RSFC correlations in the bilateral DREADD group.** Scatter plots of correlations between RSFC changes and changes of percentage of time spent in the Novel Arm in the Y-maze test, threshold at $p < 0.05$ in both CLZ-treated (orange) and CNO-treated (green) conditions. Both r-value and p-values are shown on the scatter plots.
